# 18S rDNA Sequence-Structure Phylogeny of the Trypanosomatida (Kinetoplastea, Euglenozoa) with Special Reference on *Trypanosoma*

**DOI:** 10.1101/2020.08.04.235945

**Authors:** Alyssa R. Borges, Markus Engstler, Matthias Wolf

## Abstract

**Background:** Parasites of the order Trypanosomatida are known due to their medical relevance. Trypanosomes cause African sleeping sickness and Chagas disease in South America, and *Leishmania* Ross, 1903 species mutilate and kill hundreds of thousands of people each year. However, human pathogens are very few when compared to the great diversity of trypanosomatids. Despite the progresses made in the past decades on understanding the evolution of this group of organisms, there are still many open questions which require robust phylogenetic markers to increase the resolution of trees.

**Methods:** Using two known 18S rDNA template structures (from *Trypanosoma cruzi* Chagas, 1909 and *Trypanosoma brucei* Plimmer & Bradford, 1899), individual 18S rDNA secondary structures were predicted by homology modeling. Sequences and their secondary structures, automatically encoded by a 12-letter alphabet (each nucleotide with its three structural states, paired left, paired right, unpaired), were simultaneously aligned. Sequence-structure trees were generated by neighbor joining and/or maximum likelihood.

**Results:** With a few exceptions, all nodes within a sequence-structure maximum likelihood tree of 43 representative 18S rDNA sequence-structure pairs are robustly supported (bootstrap support >75). Even a quick and easy sequence-structure neighbor-joining analysis yields accurate results and enables reconstruction and discussion of the big picture for all 240 18S rDNA sequence-structure pairs of trypanosomatids that are currently available.

**Conclusions:** We reconstructed the phylogeny of a comprehensive sampling of trypanosomes evaluated in the context of trypanosomatid diversity, demonstrating that the simultaneous use of 18S rDNA sequence and secondary structure data can reconstruct robust phylogenetic trees.

## Background

Kinetoplastids (Kinetoplastea Honigberg, 1963) are a remarkable group of unicellular organisms. They include free-living and parasite protists of invertebrates, vertebrates, and plants [1–3]. Among them, we find the obligatory parasites of the order Trypanosomatida Kent, 1880, emend Vickerman in Moreira et al. 2004 [3], including the human pathogens *T. brucei*, which causes African sleeping sickness, *T. cruzi*, the causative agent of Chagas disease in South America, and *Leishmania* species which infect and harm hundreds of thousands of people each year [2,4,5]. African trypanosomes are likely the most well-known trypanosomatids. Due to their dixenic life cycle and the extracellular lifestyle in the vertebrate blood, they have evolved interesting, and sometimes unusual, mechanisms to deal with such different environments and challenges imposed by the immune system. Thus, it does not come as a surprise that major discoveries in cell biology have been made in trypanosomes, such as antigenic variation [6], glycolysis compartmented in unique organelles [7], GPI-anchoring of membrane proteins [8], and unprecedented nucleotide modifications [9].

For harboring such a diverse group of organisms, it is unsurprising that the evolution of parasitism inside Kinetoplastea has been intriguing scientists for decades. Given that each parasitic group has closely affiliated free-living relatives and reversion to a free-living state did not occur, it is probable that at least four independent adoptions of obligate parasitism or commensalism have occurred [2,5]. Currently, the earliest diverging lineage inside Trypanosomatida is the genus *Paratrypanosoma* Votypka & Lukes, 2013, represented by one species found in mosquitoes, *Paratrypanosoma confusum* Votypka & Lukes, 2013 [10,11]. The vast majority of trypanosomatids are monoxenic parasites of insects with few dixenic genera due to the capacity of infect vertebrates, such as *Leishmania* and *Trypanosoma* Gruby, 1843 [2]. Thus, the most likely origin of *Leishmania* and *Trypanosoma* is from within monoxenic trypanosomatids, implicating that their origins were no earlier than 370 million years ago, when the invasion of land by vertebrates occurred [5,12,13]. The transmission of an insect-living trypanosomatid into a warm-blooded host has most likely occurred many times with rare successful cases [5,10,13]. So far, only *Trypanosoma* and *Leishmania* have left surviving descendants and only trypanosomes have adopted an extracellular lifestyle in vertebrates.

Once having passed the vertebrate colonization bottleneck, *Trypanosoma* radiation and adaptation to diverse vertebrate species became an unprecedented evolutionary success story. Today, these parasites prosper in essentially all vertebrate Classes, from fish to birds [13–15].

The 18S rDNA marker has been extensively used to analyze the phylogenetic relationship inside this group [5,12,16–19]. The incorporation of glycosomal glyceraldehyde phosphate dehydrogenase (gGAPDH) to the analysis allowed the generation of more resolved trees, consolidating, for example, the long-lasting question about the monophyly inside *Trypanosoma* [13,15,17–21].

The advent of modern sequencing technologies has greatly advanced our understanding of trypanosomatid phylogeny, with more new genera described in the last decade than within the past century [2,3,5]. These days, trypanosomatid phylogeny has sufficiently advanced to provide a solid framework for comparative studies, with genomic data available for more than just the medically relevant kinetoplastids. Interestingly, the basic layout of trypanosomatid genomes appears to be strikingly similar, with high overall synteny, within and between monoxenic and dixenic species [2]. The constantly growing genome data might become a powerful tool for evolutionary inference. In the future, trypanosomatids will be studied not only as infective agents of devastating neglected tropical diseases, or powerful genetic and cellular model systems, but also to unravel basic principles of the evolution of unicellular eukaryotes. What we know today is just the tip of an iceberg. The origin of *Trypanosoma*, for example, remains enigmatic. Further, the presence of prokaryotic endosymbionts and viruses in trypanosomatids, or the full biodiversity and ecological role of insect trypanosomatids remain superficially explored [2,22].

Here, we present the first large scale study in which trypanosomatid RNA secondary structure is used as an additional source of phylogenetic information. We use 18S rRNA sequence-structure data simultaneously in inferring alignments and trees. This approach was recently reviewed and shown to increase robustness and accuracy of reconstructed phylogenies [23,24]. So far, all conclusions have been made with multigene trees. The most important result of our study is that there are only a few places in our robustly supported trees where branching does not match with multigene phylogenomic trees. It is these discrepancies with which we focus our discussion, as they are potentially critical branches that are ambiguous and require more attention.

## Methods

### Taxon Sampling, Secondary Structure Prediction, Sequence–Structure Alignment, and Phylogenetic Tree Reconstruction

We have used all 240 SSU 18S ribosomal RNA gene sequences from Trypanosomatida available at NCBI (GenBank) with a sequence length >1500 nucleotides and with a full taxonomic lineage down to a complete species name. Secondary structures of the 18S rDNA sequences were obtained by homology modeling [25] using the ITS2 database [26] as well as *T. cruzi* (AF245382) and *T. brucei* (M12676) as templates (Fig S1). The two template-secondary structures (without pseudoknots) were obtained from the Comparative RNA Web (CRW) site [27]. For sequence-structure alignments, the four nucleotides multiplied by three states (unpaired, paired left and paired right) are encoded by a 12-letter alphabet [24]. Using a specific 12×12 sequence-structure scoring matrix [28], global multiple sequence-structure alignments were automatically generated in ClustalW2 1.83 [29] as implemented in 4SALE 1.7.1 [28,30]. Based on Keller et al. [23], using 12-letter encoded sequences, sequence-structure neighbor-joining (NJ) trees were determined using ProfDistS [31]. Further, using 12-letter encoded sequences, a sequence-structure maximum likelihood (ML) tree [32] - for only a representative subset of 43 sequence-structure pairs - was calculated using phangorn [33] as implemented in the statistical framework R [34]. The R script is available from the 4SALE homepage at http://4sale.bioapps.biozentrum.uni-wuerzburg.de [24]. Bootstrap support for all sequence-structure trees was estimated based on 100 pseudo-replicates. Trees were rooted with non-*Trypanosoma* sequences from Trypanosomatida.

## Results and discussion

From the analysis of 240 18S rDNA sequence-structure pairs (Fig. 1) and selection of 43 different species (Fig. 2), we obtained trees supported by high bootstraps values (> 75) on sister groups displaying the following *Trypanosoma* clades: the *Trypanosoma pestanai* Bettencourt & Franca, 1905 clade, represented in our tree by this species found in the Eurasian badger [13]; the *T. brucei* clade, consisting of trypanosome species naturally transmitted by tsetse flies, such as *Trypanosoma vivax* Ziemann, 1905, *Trypanosoma congolense* Broden, 1904, *Trypanosoma godfrey* McNamara et al., 1994, *Trypanosoma simiae* Bruce et al., 1913, *Trypanosoma equiperdum* Doflein, 1901, *Trypanosoma evansi* Steel, 1885, and *T. brucei* [13,17,19,21]; the *T. cruzi* clade, comprising mammalian trypanosomes with worldwide distribution, such as *T. cruzi, Trypanosoma rangeli* Tejera, 1920, and *Trypanosoma wauwau* Teixeira & Camargo, 2016, endemic of Latin America, *Trypanosoma conorhini* (Donovan, 1909) Shortt & Swaminath, 1928 found in Europe, South America and Africa, and *Trypanosoma dionisii* Bettencourt & Franca, 1905 distributted in Latin America, Africa, Asia and Europe [19–21,35]; the *Trypanosoma lewisi* (Kent, 1880) Laveran & Mesnil, 1901 clade, including the rodent parasites *Trypanosoma microti* Laveran & Pettit, 1910, *Trypanosoma grosi* Laveran & Pettit, 1909 and *T. lewisi* [36]; the Crocodilian clade, harboring *Trypanosoma grayi* Novy, 1906 from Africa and *Trypanosoma ralphi* Teixeira & Camargo, 2013 from South America [15,37]; the Avian clade, with *Trypanosoma corvi* Stephens & Christophers, 1908, *Trypanosoma avium* Danilewsky, 1885 and *Trypanosoma thomasbancrofti* Slapeta, 2016 [38]; the *Trypanosoma theileri* Laveran, 1902 clade, with *T. theileri*, a worldwide distributted cattle parasite, and the subclade representant *Trypanosoma cyclops* Weinman, 1972 [13,21]; and the Aquatic clade, harboring trypanosomes from fish, anurans and platypus [14,39,40]. Interestingly, the lizard/snake clade is also represented in our tree with *Trypanosoma varani* Wenyon, 1908, a snake trypanosome, branching together with the mammal parasite *Trypanosoma freitasi* Rego et al., 1957. The branching of marsupial and rodent trypanosomes inside this clade has been previously observed [36,41]. Thus, our analysis corroborates the existence of the lizard-snake/marsupial-rodent clade composed by trypanosomes transmitted by sandflies [36].

**Fig. 1:**
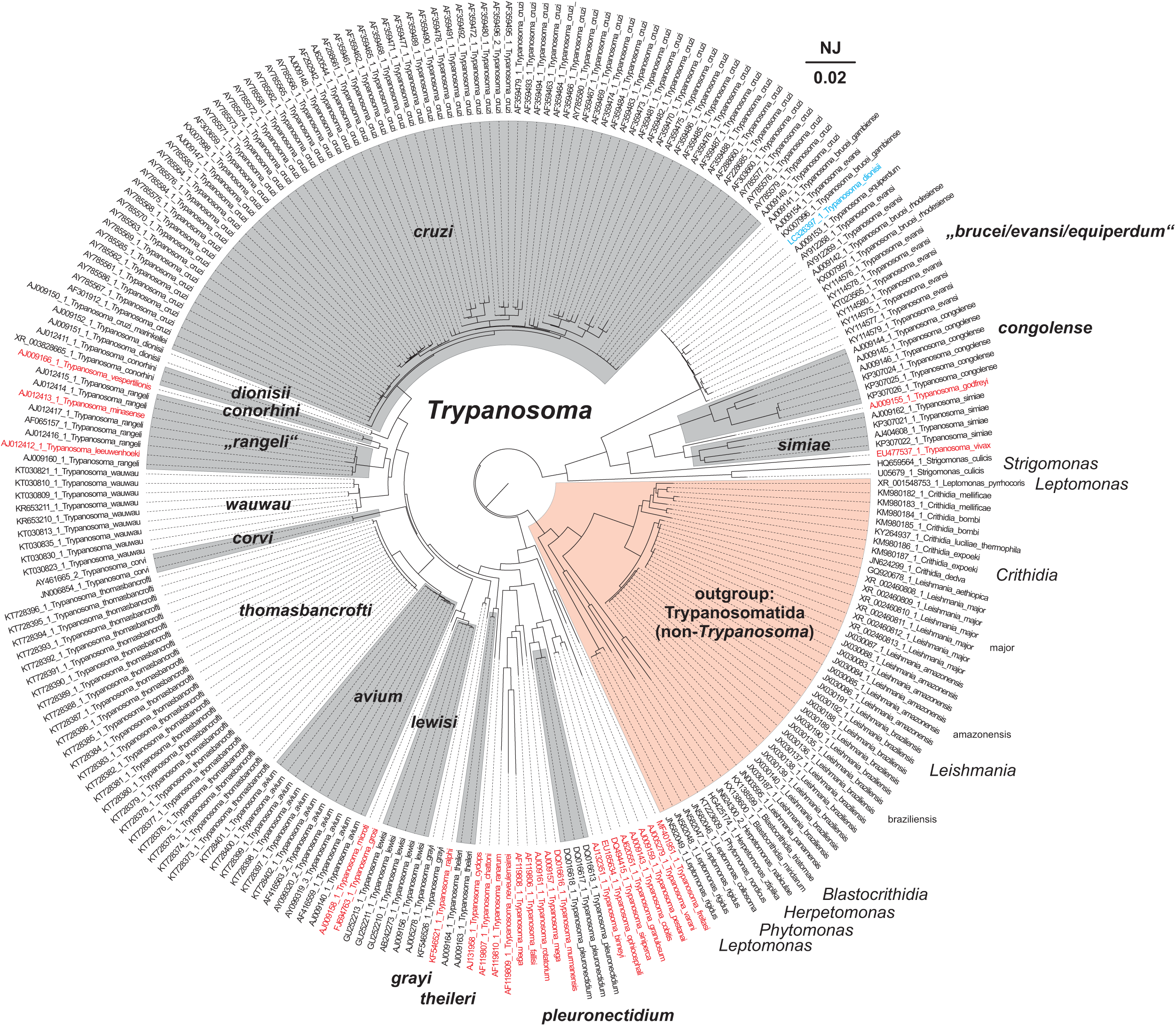
18S rDNA sequence-structure neighbor-joining (NJ) tree. obtained by ProfDistS [31]. All 240 18S rDNA sequences from Trypanosomatida (Kinetoplastea, Euglenozoa) available at NCBI (GenBank) with a sequence length >1500 nucleotides and with a full taxonomic lineage down to a complete species name have been used for the analysis. For tree reconstruction the global multiple sequence-structure alignment (.xfasta format) as derived by 4SALE [28,30] was automatically encoded by a 12-letter alphabet [24]. GenBank accession numbers accompany each taxon name. Key taxa are off and on marked in gray and additionally named alongside the tree. Non-monophyletic taxa are indicated by quotation marks. Singletons are highlighted red and polyphyletic taxa are highlighted blue. The scale bar indicates evolutionary distances. The tree is rooted at non-*Trypanosoma* sequences.

**Fig. 2:**
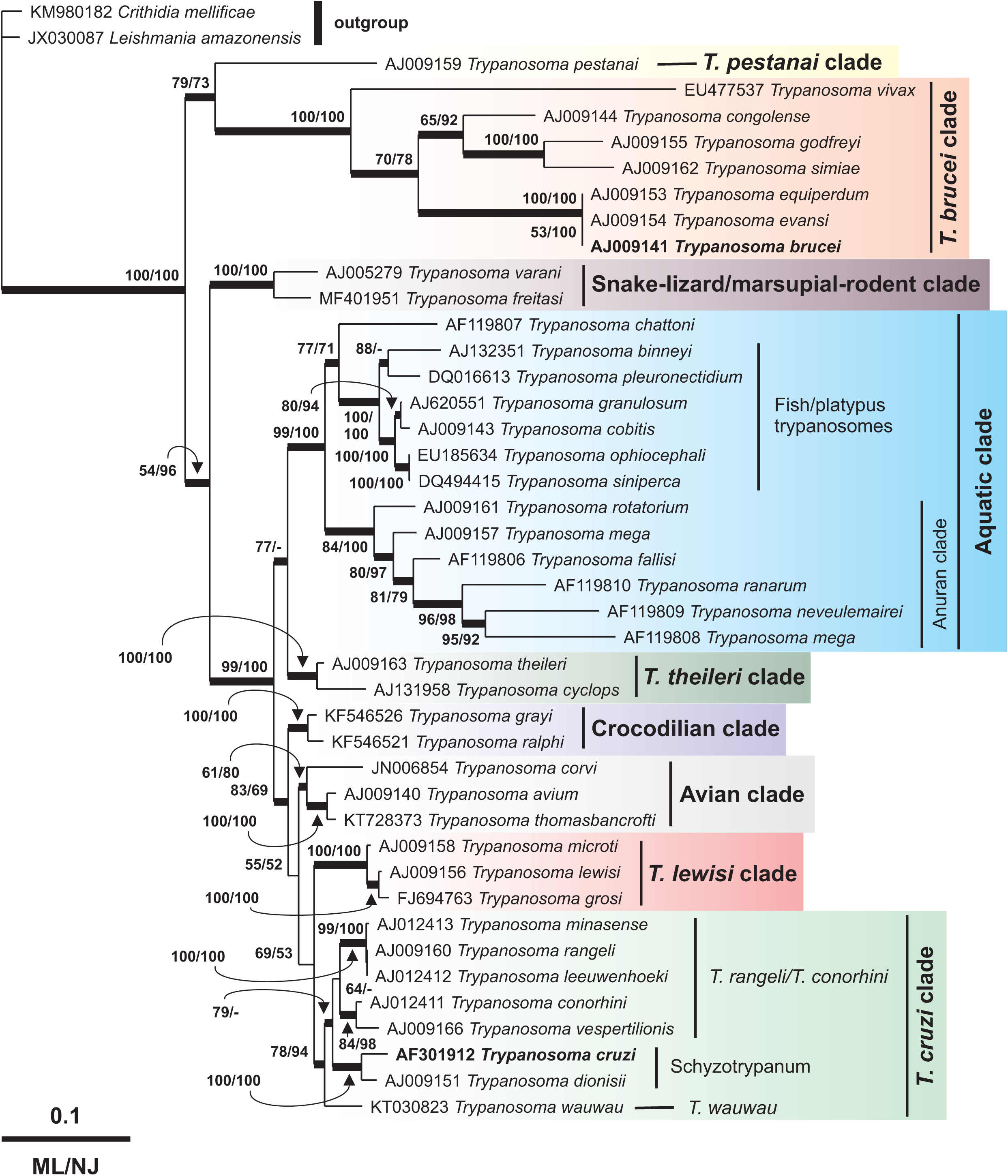
18S rDNA sequence-structure maximum likelihood (ML) tree, representative subset of 43 sequences from Fig. 1, obtained by phangorn as implemented in R [33]. Bootstrap values from 100 pseudo-replicates, mapped at the appropriate internodes, are from maximum likelihood- (ML) and neighbor-joining- (NJ, obtained by ProfDistS, Wolf et al. [31]) analyses. For NJ tree reconstruction the global multiple sequence-structure alignment (.xfasta format) as derived by 4SALE [28,30] was automatically encoded by a 12-letter alphabet [24]. For ML tree reconstruction the “one letter encoded” fasta format (12-letter alphabet) as derived by 4SALE [28,30] was used. GenBank identifiers accompany each taxon name. The scale bar indicates evolutionary distances. Highly supported branches are indicated by thicker lines. The tree is rooted at *Crithridia mellificae* (KM980182) and *Leishmania amazonensis* (JX030087). Clades discussed in the text are highlighted.

The phylogenetic analyses using sequence-structure data of 18S rDNA (Fig. 2) supports the monophyly of *Trypanosoma* as previously observed in trees constructed with partial/ complete sequences of 18S rDNA and/or gGAPDH sequences [15,17–19]. Intriguingly, in the tree obtained using a greater number of sequences (Fig. 1) *Strigomonas culicis* (Wallace & Johnson, 1961) Teixeira & Camargo, 2011 (U05679-1 and HQ659564-1) appear as a basal group of African trypanosomes (Fig. 1). To date, studies on Trypanosomatida showed a basal position of *Trypanosoma* in relation to *Strigomonas* Lwoff & Lwoff, 1931 [2,22,42]. Of notice, *S. culicis* is a monoxenic parasite of the order Diptera [22], the same order of the well-known vector of *T. brucei* clade trypanosomes, the tsetse fly. Although it may be tempting to speculate the derivation of the tsetse lifestyle of African trypanosomes from *Strigomonas*, we cannot exclude the possibility of poor sequence assembly, which could interfere with the topology observed. Finally, one sequence of the bat parasite *T. dionisii* clustered within the *T. brucei* clade (Fig. 1). This species is distributed worldwide, with its origin in Africa, and presents a high phyletic diversity [35]. However, its branching inside *T. cruzi* clade is strongly supported [13,17–19,21].

The first branching of *Trypanosoma* (Fig. 2) forms two major groups: one lineage composed by *T. brucei* and *T. pestanai* clades, and another with trypanosomes from Terrestial (*T. cruzi, T. lewisi, T. theileri*, snake-lizard/marsupial-rodent, avian and crocodilian clades) and Aquatic lineages. Thus, our tree corroborates the hypothesis of the independent evolutionary history of both human pathogens, *T. brucei* and *T. cruzi* [13]. The topology of our tree shows the Aquatic clade as a solid lineage, in accordance with previous observations [15,18,19,21]. However, the origin of this clade is still under debate. Many studies using different DNA markers, such as long (> 1.4 kb) 18S rDNA sequences, v7v8 hypervariable region of 18S rDNA and/or partial sequences of gGAPDH, showed either an early division between Aquatic and Terrestrial lineages as a single event [15,18,19,21] or in subsequent events with amphibian trypanosomes and *Trypanosoma therezieni* Brygoo, 1963 at the basis of *Trypanosoma* [17,43,44]. Interestingly, our tree suggests a later evolution of the Aquatic clade from Terrestrial trypanosomes (Fig. 2), which agrees with the insect-first hypothesis [13,36]. This hypothesis assumes that trypanosomes were originated from a monogenetic insect parasite that adapted to live inside terrestrial vertebrates and later spread to leeches and other aquatic animals, most likely through amphibians [13].

Trypanosomes of the *T. brucei* clade are virtually restricted to Africa, having an exception in *T. vivax* [45,46]. The early divergence of *T. vivax* inside the *T. brucei* clade (Fig. 1, 2) is in accordance to previous results showing a higher evolutionary rate of this species among the Salivarian trypanosomes [19,21,47]. It is interesting to consider that a previous analysis of 18S rDNA sequences revealed that members of the *T. brucei* clade show an evolutionary rate higher than other trypanosomes [47]. However, this high divergence has proven not to alter the topology of sequence-based trees [17]. Inside this clade, a low bootstrap value (ML = 53) is observed in the differentiation between *T. brucei* and *T. evansi* (Fig. 2), suggesting an unresolved positioning. In fact, the relationship between these species is controversial, with results supporting either *T. evansi* as a subspecies of *T. brucei* or showing great diversity between both depending on the *T. evansi* strains [48,49].

Regarding the other major group of our analysis (Terrestrial/Aquatic lineages), the first branching inside this group suggests the differentiation of the snake-lizard/marsupial-rodent clade as a basal group of other trypanosomes (Fig. 2). However, other studies have suggested avian trypanosomes as a basal group among terrestrial lineages [13,17]. This can be associated with the low bootstrap values of our tree in either three events: the snake-lizard/marsupial-rodent clade (ML = 54) differentiation, the divergence of crocodilian trypanosomes (ML = 55), and the internal branch of avian trypanosomes (ML = 61), which will be further explored in our discussion.

The *T. cruzi* and *T. lewisi* clades appear as sister groups in our analysis (Fig. 2), as has been previously demonstrated [17–19,21]. The *T. cruzi* clade can be subdivided into three subclades: Schyzotrypanum, *T. wauwau* (and other Neotropical bat trypanosomes, *Trypanosoma noyesi* Botero & Cooper, 2016, and *Trypanosoma livingstonei* Teixeira & Camargo, 2013) and *T. rangeli*/*T. conorhini* [21,35]. The specific sequences of *Trypanosoma minasense* Chagas, 1908 and *Trypanosoma leeuwenhoeki* Shaw, 1969 grouped with *T. rangeli* in our tree were previously considered synonyms of this species by 18S rDNA sequence analysis, explaining their positioning [13,50]. Regarding the *T. lewisi* clade, our results suggest the existence of two subclades inside the group (ML = 100), one harboring *T. microti*, and the other with *T. lewisi* and *T. grosi*. This finding is in accordance with a recent analysis of long fragments of 18S rDNA which demonstrated this subdivision despite the similarities in the v7v8 hypervariable region [51].

Concerning avian trypanosomes, our tree reflects previous findings in both the divergence between *T. avium* and *T. corvi* and the highly supported (ML = 100) proximity between *T. avium* and *T. thomasbancrofti* [38,50]. A low bootstrap value (ML = 61) is observed in the divergence of *T. corvi* and *T. avium*/ *T. thomasbancrofti*. It is interesting to note that we currently have two topologies known in literature, with the possibility of paraphyly demonstrated by analysis of long sequences of 18S rDNA [19,38]. This, however, was not observed in trees constructed with v7v8 hypervariable region of 18S rDNA, gGAPDH sequences, or concatenated trees using both [15,37,52]. Thus, our tree indicates the need for a better resolution on avian trypanosome positioning. Considering that we used only three species of avian and two species of crocodilian trypanosomes in our reconstruction, our approach represents an interesting method to be applied in further studies.

In our analysis we see the crocodilian/alligator trypanosomes (*T. grayi* and *T. ralphi*) branching together with a high support value (ML = 100) (Fig. 2). Although *T. grayi* is found in Africa and *T. ralphi* in South America [15,37,52], in a tree with distant external groups like ours, this topology is expected due to their proximity inside the Crocodilian clade [15,37]. Our tree reflects the proximity of the crocodilian trypanosomes with *T. cruzi* clade, as previously observed through full genome analysis [53]. Interestingly, crocodilian trypanosomes, such as *T. grayi* and *Trypanosoma kaiowa* Teixeira & Camargo, 2019 are tsetse-transmitted species that are not restricted to the sub-Saharan belt [15,37,53], suggesting higher adaptive plasticity of crocodilian trypanosomes.

The trypanosomes of the Aquatic lineage branched together (Fig. 2). The subgroups observed are anuran trypanosomes (*Trypanosoma rotatorium* (Mayer, 1843) Laveran, 1901, *Trypanosoma mega* Dutton & Todd, 1903, *Trypanosoma fallisi* Martin & Desser, 1990, *Trypanosoma ranarum* (Lankester, 1871) Danilewsky, 1885, and *Trypanosoma neveulemairei* Brumpt, 1928) and fish trypanosomes (*Trypanosoma siniperca* Chang, 1964, *Trypanosoma ophiocephali* Chen, 1964, *Trypanosoma cobitis* Mitrophanow, 1884, *Trypanosoma granulosum* Laveran & Mesnil, 1902, *Trypanosoma pleuronectidium* Robertson, 1906) along with the platypus parasite *Trypanosoma binneyi* Mackerras, 1959, which is in accordance to the literature [14,15,39,40]. Interestingly, the anuran parasite, *Trypanosoma chattoni* Mathis & Leger, 1911, appears in our analysis more related to fish and platypus trypanosomes than to the anuran clade. This positioning of *T. chattoni* was shown in a previous study using complete 18S rDNA sequences and non-trypanosomes as the outgroup [54]. However, recent trees using complete 18S rDNA sequences and concatenated analysis of v7v8 hypervariable region and gGAPDH rooted by other trypanosomes sustained a monophyletic anuran clade [39,40]. Thus, *T. chattoni* positioning in our tree can be related to the use of non-trypanosomes as the outgroup.

## Conclusions

18S rDNA is one of the most used markers in inferring phylogenies at higher taxonomic levels. However, this molecule has often been claimed to be inadequate for reconstructing phylogenetic relationships at lower taxonomic levels, in particular, because of its conservative rate of evolution. Thus, the use of the v7v8 hypervariable region has become popular inside the trypanosome community as a barcode suitable to describe new species along with gGAPDH [13]. However, some questions regarding the positioning of the groups still need to be answered. To this, evaluating bigger sequences and using different genetic markers can bring more information to the analysis improving resolution of the trees [13]. In this work, we demonstrate that the simultaneous use of 18S rDNA sequence and secondary structure data (i.e., the consideration of the individual secondary structures of the rRNA genes) could indeed provide important information for reconstructing more robust phylogenetic trees. Our topology highlights the need for further exploration of some groups, such as the avian and snake-lizard/marsupial-rodent clades, which are less explored in trypanosome phylogenies. Therefore, this study provides an additional fundament for upcoming phylogenetic reconstructions and/or barcoding approaches which can be used not only in trypanosomatids.

## Declarations

### Ethics approval and consent to participate

Not applicable

### Consent for publication

Not applicable

### Availability of data and materials

The sequence data analysed in this paper is publicly available in NCBI (GenBank) and their respective accession numbers are included in the trees. Data supporting the conclusions of this article are included within the article. Data and materials are available upon reasonable request to the corresponding author.

### Authors’ contributions

MW conceived the study and performed the bioinformatic analysis. ARB analyzed the data and majorly contributed to the manuscript writing. ME and MW assisted with data analysis and manuscript writing. All authors read and approved the final manuscript.

## Acknowledgments

We thank Julius Lukeš (Institute of Parasitology, Biology Centre CAS, Czech Republic) for fruitful discussions and a pre-review of our manuscript, Jaime Lisack for proof-reading, and Elisabeth Meyer-Natus for obtaining the electron microscopy image used in the graphical abstract.

## Supplementary Material

**Fig. S1:**
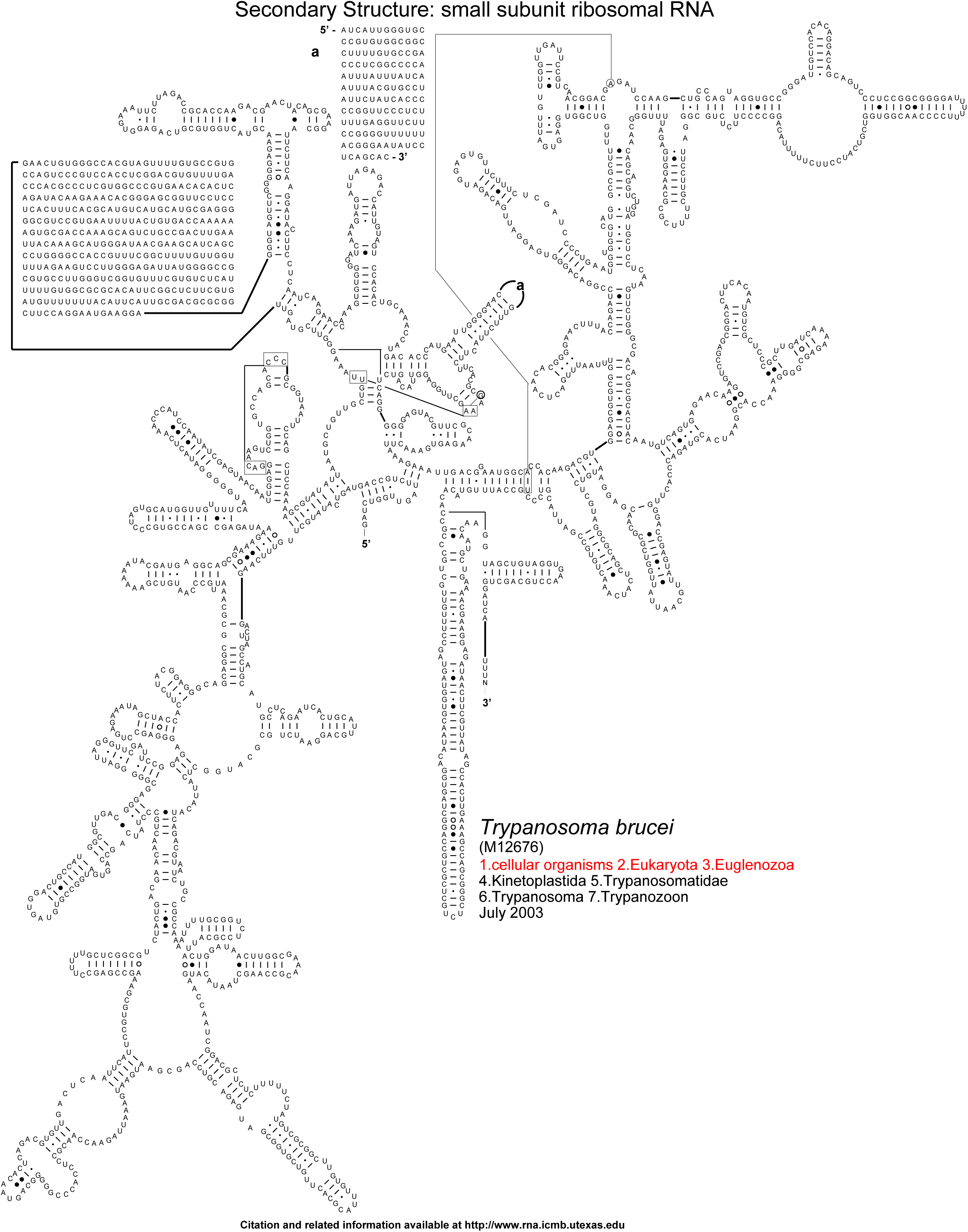
*Trypanosoma brucei* (M12676). Secondary structure: small subunit ribosomal RNA taken from the Comparative RNA Web (CRW) site [27].

